# Induction and Relief of Curiosity Elicit Parietal and Frontal Activity

**DOI:** 10.1101/195461

**Authors:** Lieke L. F. van Lieshout, Annelinde R. E. Vandenbroucke, Nils C. J. Müller, Roshan Cools, Floris P. de Lange

**Affiliations:** Donders Institute for Brain, Cognition and Behaviour, Radboud University, P.O. Box 9101, 6500 HB Nijmegen, The Netherlands.; Department of Psychiatry, Radboud University Medical Centre, P.O. Box 9101, 6500 HB Nijmegen, The Netherlands.; Helen Wills Neuroscience Institute, University of California, Berkeley, California 94720-3192, USA.; Donders Institute for Brain, Cognition, and Behaviour, Radboud University Medical Centre, 6525 EN Nijmegen, the Netherlands.

## Abstract

Curiosity is a basic biological drive, but little is known about its behavioral and neural mechanisms. We can be curious about several types of information. On the one hand, curiosity is a function of the expected value of information, serving primarily to help us maximize reward. On the other hand, curiosity can be a function of the uncertainty of information, helping us to update what we know. In the current studies, we aimed to disentangle the contribution of information uncertainty and expected value of rewards to curiosity in humans of either sex. To this end, we designed a lottery task in which uncertainty and expected value of trial outcomes were manipulated independently, and examined how neural activity and behavioral measures of curiosity were modulated by these factors. Curiosity increased linearly with increased outcome uncertainty, both when curiosity was explicitly probed as well as when it was implicitly tested by people’s willingness to wait. Increased expected value, however, did not strongly relate to these curiosity measures. Neuroimaging results showed greater BOLD response with increasing outcome uncertainty in parietal cortex at the time of curiosity induction. Outcome updating when curiosity was relieved resulted in an increased signal in the insula, orbitofrontal cortex and parietal cortex. Furthermore, the insula showed a linear increase corresponding to the size of the information update. These results suggest that curiosity is monotonically related to the uncertainty about one’s current world model, the induction and relief of which are associated with activity in parietal and insular cortices respectively.

**Significance statement:** Humans are curious by nature. When you hear your phone beep, you probably feel the urge to check the message right away, even though the message itself likely doesn’t give you a direct reward. In this study, we demonstrated that curiosity can be driven by outcome uncertainty, irrespective of reward. The induction of curiosity was accompanied by increased activity in the parietal cortex, whereas the information update at the time of curiosity relief was associated with activity in insular cortex. These findings advance our understanding of the behavioral and neural underpinnings of curiosity, which lies at the core of human information-seeking and serves to optimize the individual’s current world-model.

## Introduction

In daily life, we receive enormous amounts of information from our surroundings. Curiosity relating to the content and use of some of this information is a basic biological drive, but little is known about the underlying behavioral and neural mechanisms (Gottlieb et al., 2013; Kidd & Hayden, 2015). An obvious reason to be curious about information is to reach a goal or to maximize rewards. In these cases, curiosity is instrumental for obtaining reward and increases as a function of the expected value of information (Daw et al., 2006).

However, sometimes information can be appealing in itself. Even animals are particularly curious when information provides a substantial update of what they know (Blanchard et al., 2015; Bromberg-Martin & Hikosaka, 2009, 2011). For instance, monkeys performing an information-seeking choice task choose to have outcomes of risky gambles revealed immediately, instead of remaining in a state of uncertainty when waiting for the outcome (Bromberg-Martin & Hikosaka, 2009, 2011). Interestingly enough, this is the case even though receiving information is not instrumental, i.e. will not help them to improve performance or to obtain higher rewards. In fact, monkeys are willing to sacrifice a substantial amount (20% - 33%) of primary reward to get advance information (Blanchard et al., 2015). Some researchers (Dinsmoor, 1983; Shannon & Weaver, 1949) have highlighted this phenomenon as an exemplary departure from normatively optimal (reward-guided) behavior (but see Beierholm & Dayan, 2010, for an alternative account).

Here, we aimed to elucidate behavioral and neural determinants of curiosity in humans. We designed a lottery task in which two potential sources of curiosity are manipulated independently: outcome uncertainty and expected value of the outcome. Consistent with previous experiments on observing behavior (Bromberg-Martin & Hikosaka, 2009, 2011), observing the outcome was designed to not have any impact on received rewards. As such, this paradigm allowed us to disentangle uncertainty from value accounts of observing behavior.

On each trial, participants received a lottery with an uncertain outcome and associated monetary reward (Figure 1). We probed participants’ curiosity on each trial either explicitly (Experiment 1, 3) or implicitly (Experiment 2) by investigating participants’ willingness to wait for the outcome (Kang et al., 2009; Marvin & Shohamy, 2016). We hypothesized that participants are particularly curious in situations of high uncertainty, because information then leads to a large belief update (information prediction error). Using functional magnetic resonance imaging (fMRI) we examined which brain areas were modulated by outcome uncertainty and expected value upon lottery presentation, and their updates upon outcome presentation. We hypothesized that during lottery presentation, brain regions associated with uncertainty or risk, such as insular cortex (Critchley et al., 2001; Jepma et al., 2012; Paulus et al., 2003; Preuschoff et al., 2006, 2008) and parietal cortex (Foley et al., 2017; Huettel et al., 2005), would be modulated by outcome uncertainty. During outcome presentation, we expected increased activity in brain regions involved in curiosity relief and information updating, such as the insula (Jepma et al., 2012; Preuschoff et al., 2008), orbitofrontal cortex (Blanchard et al., 2015; Jepma et al., 2012) and ventral striatum (Jepma et al., 2012; Wittmann et al., 2008). In addition, we expected activity in reward-related areas (e.g. ventral striatum) to be modulated as a function of expected value (Knutson et al., 2005; O’Doherty, 2004) and reward prediction error (Bayer & Glimcher, 2005; Daw & Doya, 2006; Schultz et al., 1997).

**Figure 1.**
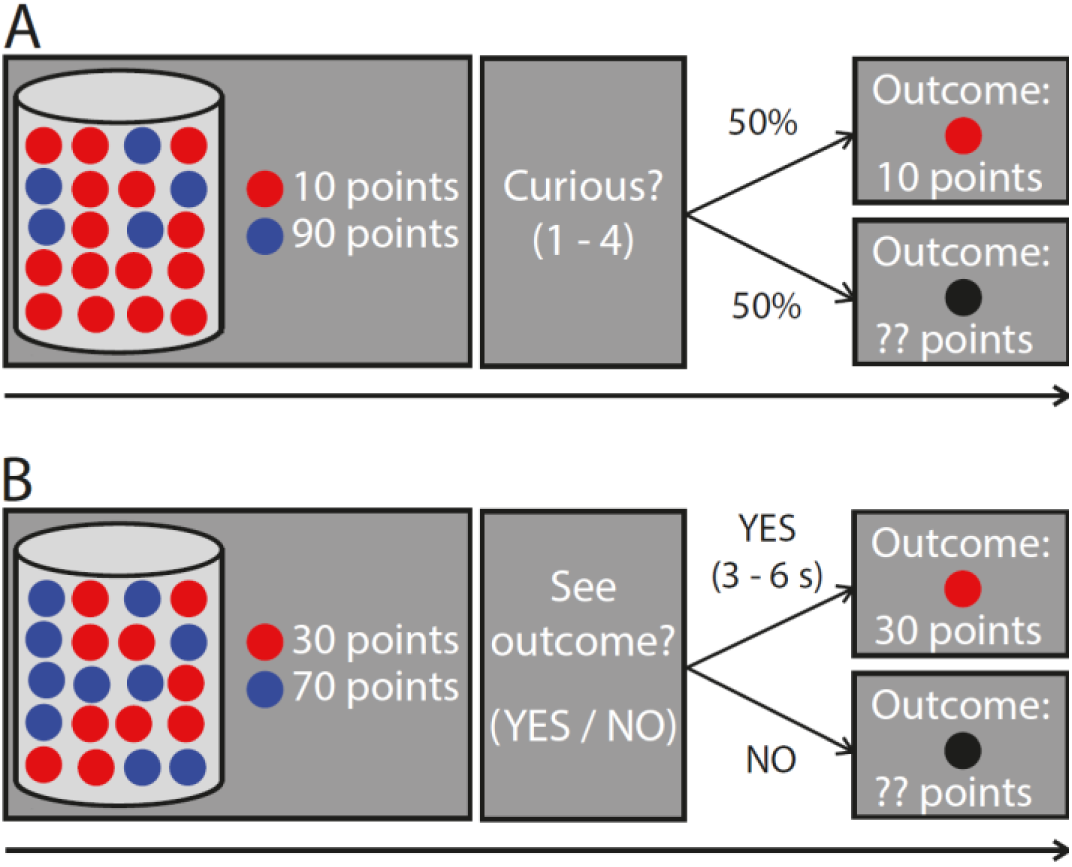
(A) Schematic figure of Experiment 1 and Experiment 3.Participants saw a screen on which a vase with 20 marbles was depicted, either of which could be red or blue, and the points associated with these marbles. Next, participants indicated how curious they were about seeing the outcome of the vase (1 – 4). There was a 50% chance of seeing the outcome, regardless of the participants’ curiosity response. See text for details on the timing of the experiments (for Experiment 1 see *Procedure – Behavioral* and for Experiment 3 see *Procedure - fMRI*). (B) Schematic figure of Experiment 2. The task was similar to Experiment 1 and 3, except that participants indicated whether they wanted to see the outcome of a trial or not. If they responded “Yes”, they had to wait 3 – 6 seconds before the outcome was presented to them and if they responded “No” the outcome was not presented to them. See text (*Procedure – Behavioral*) for details on the timing of the experiment.

To preview, we found robust increases in curiosity with increasing outcome uncertainty, whereas expected value did not robustly modulate curiosity. Outcome uncertainty generated activity in parietal cortex, whereas its relief was associated with increased activity in a network comprising the insula, orbitofrontal cortex and parietal cortex. Furthermore, we found a parametric increase in insular activity with increasing information prediction error. Together, these findings suggest that curiosity can be conceptualized as a desire to improve one’s current world model and identify neural effects that accompany this desire.

## Materials and Methods

### Participants

Twenty-four healthy individuals participated in Experiment 1, in which we explicitly probed curiosity (17 women, age 23.1 ± 3.8, mean ± SD). Another twenty-five healthy individuals participated in Experiment 2, in which we implicitly probed curiosity by investigating participants’ willingness to wait. One participant was excluded due to a lack of variation in responding (chose to wait in > 98% of all trials). Therefore, the final sample of Experiment 2 consisted of twenty-four participants (17 women, age 24.6 ± 5.5, mean ± SD). Finally, twenty-eight healthy individuals participated in Experiment 3, in which we explicitly probed curiosity while non-invasively recording neural activity using fMRI. All participants were right-handed, screened for MR compatibility and had normal or corrected-to-normal vision. Two participants were excluded due to a technical problem of the MRI scanner. One participant was excluded because of too many missed trials (> 10% of all trials) and one participant was excluded due to self-reported difficulties viewing the screen. The final sample for Experiment 3 consisted of 24 participants (18 women, age 24.1 ± 3.2, mean ± SD). All three experiments were approved by the local ethics committee (CMO Arnhem-Nijmegen, The Netherlands) under the general ethics approval (“Imaging Human Cognition”, CMO 2014/288) and the experiments were conducted in compliance with these guidelines. All participants gave written informed consent according to the declaration of Helsinki prior to participation.

### Procedure - Behavioral

*Experiment 1* – Each trial started with an image of a vase containing twenty marbles (Figure 1A). Each of the marbles could be either red or blue. In total, four vase configurations were possible: (1) 100% vases: all marbles had the same color, (2) 95%-5% vases: 19 marbles had one color and 1 marble had the other color, (3) 75%-25% vases: 15 marbles had one color and 5 marbles the other color, (4) 50-50% vases: 10 marbles had one color and 10 marbles the other color. Both colored marbles were associated with points participants could earn. The points for each of the marbles varied between 10 and 90 (in steps of 10). All combinations of points associated with red and blue marbles were possible. The participants were informed that on each trial, one marble would be selected from the vase and that they would be awarded with the points associated with this marble. The first screen, on which the vase, the marbles and the points associated with the marbles were depicted, was presented for 3000 ms. Then a blank screen was presented for 500 ms, followed by a response screen during which participants could indicate how curious they were about seeing the outcome of that trial (“How curious are you about the outcome?”). The curiosity scale ranged from 1 – 4. The response screen was presented until the participant responded, with a response limit of 4000 ms. The response screen was followed by a blank screen (500 ms) and an outcome screen (2000 ms). On each trial, participants had a 50% chance that their curiosity would be satisfied by seeing the outcome (curiosity relief) and a 50% chance that the outcome was withheld. This manipulation was explicitly instructed to subjects and it uncoupled curiosity responses from the actual receipt of the outcome (thus rendering Pavlovian bias accounts of our observing behavior less likely, Beierholm and Dayan, 2010). In the follow-up fMRI experiment (Experiment 3, see below) it enabled us to investigate the neural consequences of curiosity relief (Curiosity Relief – Yes versus Curiosity Relief – No). The outcome screen depicted the vase, the marbles and points associated with the marbles again, together with a box in which they saw the colored marble that was selected and how many points they earned. When the outcome was not presented, participants saw a black marble instead of a colored marble and question marks at the location of the number of points. This way, the amount of visual input was roughly comparable between presented and not presented outcomes. After a trial ended, there was a blank screen with a jittered duration between 1000 and 2000 ms (uniformly distributed). Importantly, participants had no way of influencing whether they would observe the outcome of a particular trial, or what the outcome of that trial would be. However, participants knew that a marble would be selected in every trial and that they would be awarded with the points associated with that marble, even if the outcome was not presented. They were told that the total amount of points they earned in total would be converted to a monetary bonus at the end of the experiment.

The trials were pseudo-randomized such that the same vase configuration was never presented more than 4 trials in a row. Each vase configuration was presented on 144 occasions, except for the 100% vases which were only presented 45 times. These 100% trials were included as a control to check participants’ compliance to the task; we expected people not to be curious about trials of which they already knew the outcome. In total, the participants completed 477 trials, divided in 9 blocks of 53 trials. After each block, the participants were instructed to take a short break if they wanted. The experiment lasted ∼ 75 minutes in total.

*Experiment 2 –* In Experiment 2 we aimed to investigate participants’ curiosity more implicitly by means of testing their willingness to wait to see the outcome. We used willingness to wait because it is a well-established measure of the motivational value of an item (Frederick et al., 2002), which has been previously linked to curiosity (Kang et al., 2009; Marvin & Shohamy, 2016). The trial setup for Experiment 2 was similar to Experiment 1 (Figure 1B). The start of the trial was identical, but instead of giving a curiosity response, participants indicated whether they wanted to see the outcome of that trial (“Do you want to see the outcome?”) by pressing either “Yes” or “No”. If they pressed “Yes”, they saw a blank screen with a jittered duration between 3000 and 6000 ms (uniformly distributed) before they saw a screen on which the outcome was presented (2000 ms). If they pressed “No”, a blank screen was presented briefly (500 ms), followed by a screen on which the outcome was not presented (2000 ms). The outcome screens looked identical to the outcome screens in Experiment 1. The next trial started after a jittered inter-trial interval between 1000 and 2000 ms (uniformly distributed).

The trials were again pseudo-randomized in a way that the same vase configuration was never presented in more than 4 trials in a row. The total willingness to wait experiment consisted of 261 trials. Each vase configuration was presented on 72 occasions except for the 100% trials, which were only presented 45 times. The total duration of the experiment depended on for how many trials the participants indicated that they were willing to wait to see the outcome. Again, participants were told that that a marble would be selected on every trial and that they would be awarded with the points associated with that marble, also when they decided not to wait to see the outcome. The total amount of points they earned would be converted to a monetary bonus at the end of the experiment.

### Experimental Design and Statistical Analysis - Behavioral

*Experiment 1* – The behavioral analyses of Experiment 1 were performed using MATLAB (The MathWorks, RRID:SCR_001622) and SPSS (RRID:SCR_002865). We investigated whether there was a relationship between outcome uncertainty (OU) and the curiosity ratings, as well as between expected value (EV) and the curiosity ratings. In order to do so, a value of OU was calculated for each trial by multiplying the Shannon Entropy by the absolute difference between the red (*x*_1_) and blue (*x*_2_) marble points:

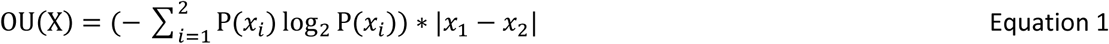

where P(*x*_*i*_) denotes the probability that a marble (i) would be drawn. Thereby, OU reflected a combination of two types of variance: the variance in the division of the marbles (as indicated by the Shannon Entropy) and the variance in points associated with both marbles (indicated by the absolute difference between the marble points). In turn, EV values were calculated (Equation 2) by the sum of the probability that a red marble would be drawn (*p*_1_) multiplied by the number of points associated with the red marble (*x*_1_) and the probability that a blue marble would be drawn (*p*_2_) multiplied by the number of points associated with the blue marble (*x*_2_):

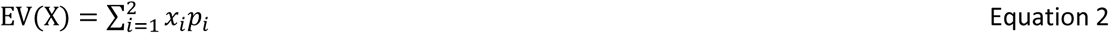

We investigated whether there was a relationship between OU and/or EV and the curiosity ratings using a Univariate General Linear Model (GLM) with dependent variable “Curiosity Rating (1 – 4)” and random factors “Participant”, “OU” and “EV”.

*Experiment 2* – For Experiment 2, we investigated whether there was a relationship between OU and/or EV and whether participants were willing to wait to see the outcome of a trial. These data were analyzed using a binomial logistic regression with dependent variable “Willingness to wait (Yes/No)” and independent variables “Participant”, “OU” and “EV”. Statistical significance of the model was assessed using χ^2^ and the amount of variance explained by the model was estimated using Nagelkerke R^2^. The values for odds ratio (OR) for each independent variable were used to determine the directionality of any significant effects.

### Data visualization – Behavioral

*Experiment 1* – To visualize the behavioral data, the values of OU and EV were divided in percentile bins, such that the 10^th^ percentile represents the 10% lowest values of either OU or EV, the 20^th^ percentile represents the 10% - 20% of the lowest values, etc. This was done to enable us to visually compare the effects of OU and EV. To plot the data, we performed the same analysis on the binned data as described above (see *Experimental Design and Statistical Analysis – Behavioral: Experiment 1).* However, to isolate the contributions of OU and EV to curiosity, we added “Participant” and “EV” as covariates to visualize the effects of “OU” on curiosity. The same analysis was performed to visualize the effects of “EV” on curiosity, but this time “Participant” and “OU” were added as covariates. This provided a clear view of the independent effects of OU and EV, because even though OU and EV were manipulated independently and showed no linear correlation with each other, they did show a quadratic correlation such that middle values of OU were associated with higher values of EV. These analyses were performed for each participant separately and the mean curiosity scores for each percentile were calculated by averaging over participants.

*Experiment 2* – To visualize the data for the willingness to wait study, the values of OU and EV were divided in percentile bins as described for Experiment 1. To plot the data, we used a Univariate General Linear Model (GLM) with dependent variable “% Willingness to wait”, random factor “OU” and covariates “Participant” and “EV”. The same analysis was performed to visualize the relationship between EV and willingness to wait, but this time “EV” was added as random factor and “Participant” and “OU” as covariates. Again, these analyses were performed for each participant separately and the mean willingness to wait scores for each percentile were calculated by averaging over participants.

### Procedure - fMRI

*Experiment 3* - The trial setup for the fMRI study (Experiment 3) was similar to the setup of Experiment 1 (Figure 1A), except that the timing was adjusted to allow for analyses based on the BOLD response. First, participants saw a screen on which the vase containing the marbles and the points associated with the marbles was presented (4000 ms). Next, participants saw a screen during which they could indicate their curiosity for the outcome on a scale from 1 – 4. This screen was presented for 2500 ms. Then there was a blank screen presented for a jittered duration between 2500 and 4500 ms (uniformly distributed). After the blank screen, participants saw a screen on which the outcome of a trial was either presented or not presented (50% chance) and this screen was displayed for 2000 ms. When the outcome was presented, they saw the colored marble in the middle of the screen and below that the number of points the participant earned. When the outcome was not presented, they saw a black marble in the middle of the screen with question marks at the location where otherwise the number of points would have been presented. Then there was an inter-trial interval consisting of a blank screen which was presented for a jittered duration between 3500 and 4500 ms (uniformly distributed). After every 9 trials, the duration of the blank screen was prolonged to be jittered between 9500 and 10500 ms (uniformly distributed). From these prolonged blank screens, a baseline was estimated.

Moreover, only 75%-25% and 50%-50% vases were used to reduce differences in visual processing between the different vase configurations. The participants completed a total of 180 trials divided in 4 sessions of 45 trials. After each session, the participants were able to take a short break. In total, the experiment lasted ∼ 60 minutes. Also for this experiment, participants knew that they had no way of influencing whether they would observe the outcome of a trial or not and what that outcome would be. Again, a marble would be selected in every trial and all the points would be added together and converted to a monetary bonus at the end of the experiment. After functional image acquisition, an anatomical image was acquired.

### fMRI acquisition and preprocessing

Functional images were acquired using a multiband imaging sequence (TR = 769 ms, TE= 39 ms, 54 transversal slices, voxel size of 2.4 x 2.4 x 2.4 mm, multiband acceleration factor 6, 52° flip angle). Using the AutoAlign head software by Siemens, we ensured a similar FOV tilt across participants. Anatomical images were acquired using a T1-weigted MP-RAGE sequence, using a GRAPPA acceleration factor of 2 (TR = 2300 ms, TE = 3.03 ms, voxel size 1 x 1 x 1 mm, 192 transversal slices, 8° flip angle).

fMRI data preprocessing was carried out using FSL (RRID:SCR_002823) and SPM8 (RRID: SCR_007037). The first seven volumes of each run were discarded to correct for T1 equilibration. The following pre-statistics processing was applied; motion correction using MCFLIRT (Jenkinson et al., 2002); non-brain removal using BET (Smith, 2002); spatial smoothing using a Gaussian kernel of FWHM 6.0 mm; grand-mean intensity normalisation of the entire 4D dataset by a single multiplicative factor. Registration to high resolution structural images was carried out using FLIRT (Jenkinson et al., 2002; Jenkinson & Smith, 2001). Registration from high resolution structural to standard space was then further refined using FNIRT nonlinear registration (Andersson et al., 2007a, 2007b). Next, ICA-AROMA (Pruim et al., 2015) was used to reduce motion-induced signal variations in the fMRI data after which the data were high-pass filtered at 1/100 Hz. Finally, data were normalized to standard space and further analysed using SPM8.

### Experimental Design and Statistical Analysis - fMRI

*fMRI behavioral analyses* - For the behavioral data of Experiment 3, the same analysis was performed as for Experiment 1 (see *Experimental Design and Statistical Analysis - Behavioral*) using MATLAB (The MathWorks, RRID:SCR_001622) and SPSS (RRID:SCR_002865). Behavioral data visualization for Experiment 3 was done in the same way as for Experiment 1 (see *Data Visualization – Behavioral*).

*fMRI BOLD analyses* - For each subject, data was modeled using an event-related GLM. The first regressor modeled the period when the vase was presented, serving to induce curiosity in the participants (Curiosity Induction), and had a duration of 4000 ms. The second and third regressor modeled the moment when the outcome of a trial was either presented or not presented and thereby relieved (or not) curiosity in the participants (Curiosity Relief – Yes / No). Both regressors had a duration of 2000 ms. The fourth regressor represented the moment the participants pressed a button to indicate their level of curiosity (Button press) and was modeled as a stick function. The fifth regressor was the baseline (Baseline), which started after every 9 trials when the prolonged blank screen was presented. The duration of this regressor was the duration that the blank screen was presented. All regressors were convolved with a canonical haemodynamic response function (HRF; Friston et al., 1998).

For each trial, values for OU and EV were calculated (see *Experimental Design and Statistical Analysis – Behavioral*) and simultaneously included as a parametric modulations of activity during the Curiosity Induction period in the GLM. To this end, the unit height HRF of the Curiosity Induction regressor was convolved with vectors of parametric weights that reflected the trial-by-trial fluctuations of OU and EV. OU and EV were used as parametric modulators at the moment of Curiosity Induction, because this was the moment when participants process the vase and when they could make an estimation of the uncertainty of the outcome of a trial (OU) and how much they will approximately earn in that trial (EV).

Furthermore, at the moment of trial outcome (Curiosity Relief), we looked at which brain areas respond more strongly to receiving information about the outcome than to not receiving this information and vice versa. We did so by investigating which brain areas were active for the Curiosity Relief contrast (Curiosity Relief – Yes > Curiosity Relief – No) and which brain areas were active for the opposite contrast. We used a primary voxel threshold of *p* < .001 (uncorrected). Inference was based on a cluster-level correction of *p* < .05 (Family-Wise Error, FWE).

*ROI analyses* - Additional to the whole brain fMRI analyses, we defined regions of interest (ROIs) based on their significant overall activity modulation during the Curiosity Relief period, using the SPM toolbox MarsBaR (Brett et al., 2002; RRID:SCR_009605). The rationale for selecting ROIs based on the Curiosity Relief contrast (Curiosity Relief – Yes > Curiosity Relief – No) was that these regions responded more strongly to receiving information about the outcome than to not receiving this information, indicating that they might play a role in processing information about the outcome. We aimed to investigate whether these regions play a role in processing the extent to which participants received an update of information (information prediction error) and reward (reward prediction error). The ROIs based on the opposite contrast (Curiosity Relief – No > Curiosity Relief – Yes) respond stronger to not receiving information about the outcome compared with receiving this information. We aimed to investigate whether these regions play a role in processing the extent to which participants were left with uncertainty about what the outcome would have been (which we refer to as negative information prediction error, see below).

To investigate whether there was a modulation of activity as a function of information and/or reward prediction error effects in these ROIs, we calculated values for the reward prediction error (RPE), the information prediction error when the outcome was presented (IPE_relief_) and the information prediction error when the outcome was not presented (IPE_no relief_). The RPE (Equation 3) is the discrepancy between the number of points the participant received (x_shown_) on a given trial (X) and the number of points the participant expected to receive (EV). The RPE was positive when the participant received more points than expected and negative when the participant received less points than expected. Note that when the outcome was not presented, there was no value for RPE.

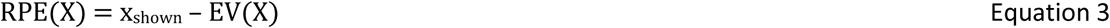

Besides information about the reward, participants get an update of information when the outcome was presented (when they know which marble was selected for them; IPE_relief_; Equation 4).

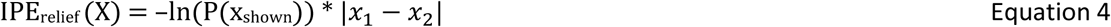

IPE_relief_ reflects the amount of information that the participant received when being presented with the outcome of a trial.It is calculated by means of the negative natural log of the probability that the shown outcome would be presented P(x_shown_).This value is than multiplied by the absolute difference between the red (x_1_) and blue (x_2_) marble points. On the contrary, when the outcome is not presented, participants get no update of information and they remain uncertain about which marble was selected for them (IPE_no relief_). Therefore, the IPE_no relief_ was calculated as a function of OU (i.e. the more uncertain participants were at the beginning of a trial, the more uncertainty will remain after not seeing the outcome; Equation 5). Here, OU(X) reflects the outcome uncertainty of a given trial X (Equation 1).

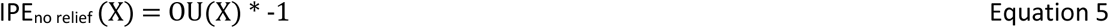

Note that the value of IPE_relief_ was always positive and the value of IPE_no relief_ was always negative. In other words: when the outcome was presented, IPE was positive and when the outcome was not presented IPE was negative.

The values of RPE, IPE_relief_ and IPE_no relief_ were simultaneously included as parametric weights for Curiosity Relief in the GLM. The unit height of “Curiosity Relief - Yes” was convolved with vectors of parametric weights that reflected the trial-by-trial fluctuations of IPE_relief_ and RPE. The same was done for “Curiosity Relief - No”, but then with a vector of parametric weights reflecting the fluctuations of IPE_no relief_. Note again that no parametric weights for RPE could be calculated when no outcome is presented, since there is no update about what reward participants received.

For each of the significant ROIs found for the Curiosity Relief contrast, we tested the parameters for RPE, IPE_relief_ and IPE_no relief_ against 0 using one-sample t-tests. The same was done for ROIs that were selected based on the opposite contrast (Curiosity Relief – No > Curiosity Relief - Yes).

Finally, given our specific hypothesis specified in the introduction, that reward-related areas (e.g. ventral striatum) would be active as a function of expected value, reward prediction error and possibly also as a function of information updating, we investigated mean extracted signal from an a priori defined ROI in the ventral striatum (with MarsBaR, Brett et al., 2002; RRID:SCR_009605). This ROI was selected from a prior resting state MRI study in which the striatum was parcellated into (anatomically plausible) subregions in a functional data-driven manner, based on intrastriatal functional connectivity analyses that correlated each striatal voxel with all other striatal voxels (Piray et al., 2015). Within the ventral striatum, we investigated the parametric modulation effects of OU, EV, RPE, IPE_relief_ and IPE_no relief_. We tested the parameters for these contrasts against 0 using one-sample t-tests.

## Results

### Behavioral results

*Experiment 1* – The aim of Experiment 1 was to investigate whether and how participants’ curiosity was modulated by outcome uncertainty (OU) and expected value (EV).

Curiosity strongly and monotonically increased with increasing OU (*F*_1,15_ = 103.07, *p* <.001). Conversely, there was no significant modulation of curiosity by EV (*F*_1,87_ = 1.25, *p* = .15; Figure 2A). There was an interaction between OU and EV (*F*_1,68_ = 1.91, *p* < .001), such that the relationship between OU and curiosity was stronger for medium values of EV, compared with high and low values. This is likely due to the design of the experiment, which had a restricted range of OU values for extreme values of EV, such that extreme values of EV were always associated with low values of OU.

**Figure 2.**
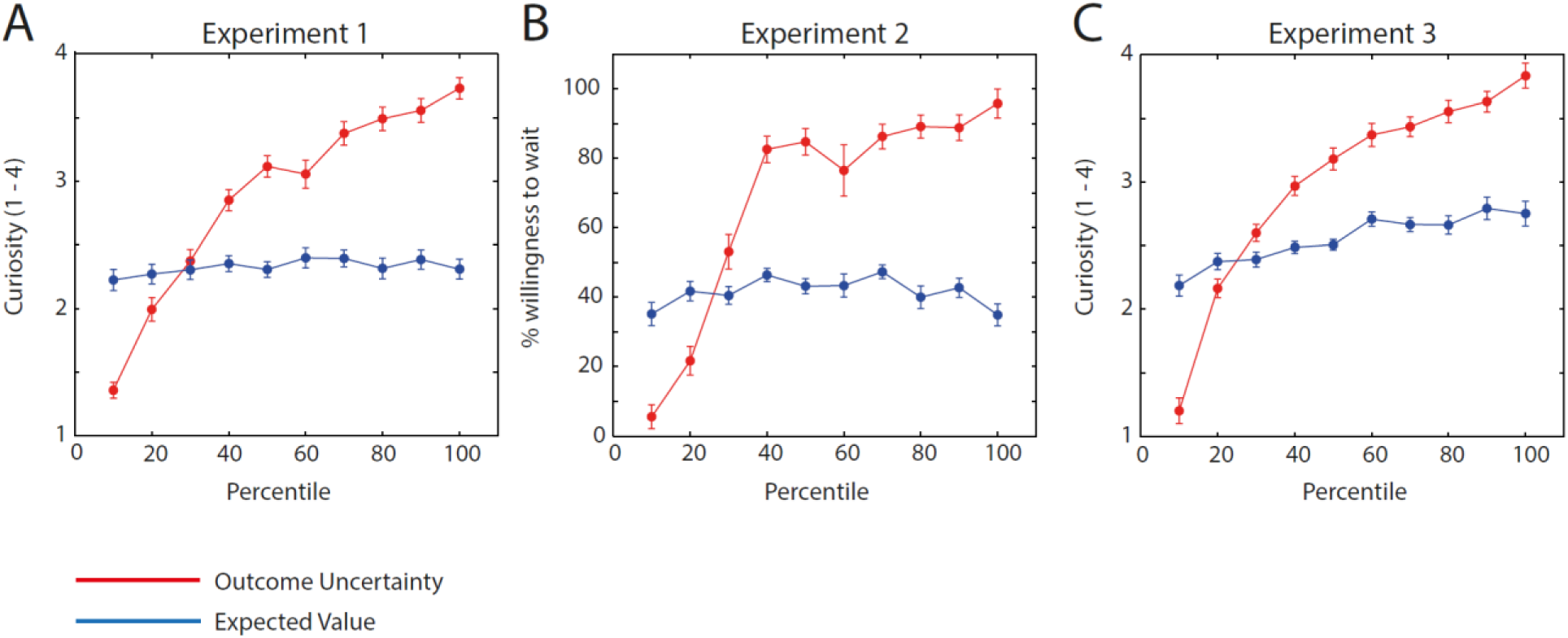
Behavioral results of the three experiments. The x-axis depicts percentile bins of the values of OU (in red) and EV (in blue). The y-axis depicts the mean curiosity rating (A and C) or the percentage willingness to wait (B) for each percentile of OU and EV. In all panels, the effects of EV on curiosity are controlled for OU, and the effects of OU on curiosity are controlled for EV. For details on behavioral data visualisation see *Data visualisation – Behavioral*. Error bars depict the standard error of the mean (SEM).

*Experiment 2 –* In Experiment 2, we investigated whether participants would be more willing to wait to see the outcomes of trials they indicated to be more curious about in the first experiment. In other words: were people willing to sacrifice time in order to satisfy their curiosity?

The results show that people were more willing to wait (χ^2^(3) = 2944.29, *p* < .001, Nagelkerke pseudo R^2^ = .505) when OU was higher (Odd’s Ratio = 1.11, *p* < .001), but that willingness to wait was not modulated by EV (Odd’s Ratio = 1.00, *p* = .98). These results are consistent with Experiment 1, showing that people not only indicated to be more curious when OU is higher, but that they were also more willing to wait to see this outcome (Figure 2B). Similarly, people were not more willing to wait for trials with high compared with low EV, supporting the findings of Experiment 1. In Experiment 1, there was a significant interaction effect between OU and EV on curiosity. Therefore, we investigated whether adding the interaction in this model would improve the model-fit. However, the model did not get better by adding the interaction between OU and EV (χ^2^(4) = 2.48, *p* = .12). This indicates that the effects of OU on willingness to wait did not differ between different values of EV.

*Experiment 3 –* As in Experiment 1, curiosity increased with increasing OU (*F*_1,14_ = 69.87, *p* < .001), replicating the finding of Experiment 1. In Experiment 3, however, curiosity also increased with increasing EV (*F*_1,30_ = 3.26, *p* < .001; Figure 2C), although the magnitude of the modulation was markedly smaller. There was also an interaction between OU and EV (*F*_1,61_ = 3.00, *p* < .001), such that the relationship between OU and curiosity was stronger for medium values of EV, compared with high and low values.

### Neuroimaging results

*Curiosity Induction* - First, we examined activity when curiosity was induced (during vase presentation) as a function of OU and EV. There was one cluster that showed a significant increase in activity with increasing OU, located in the left inferior parietal lobule (Table 1, see Appendix; Figure 3). Somewhat surprisingly, there was a large network of regions that showed a significant increase in activity with decreasing OU (Table 1, see Appendix; Figure 3), including the hippocampus, precuneus and several clusters within the temporal and frontal lobe. No significant positive modulation of EV was observed, while activity in the calcarine sulcus increased as a function of decreased EV (Table 1, see Appendix).

**Figure 3.**
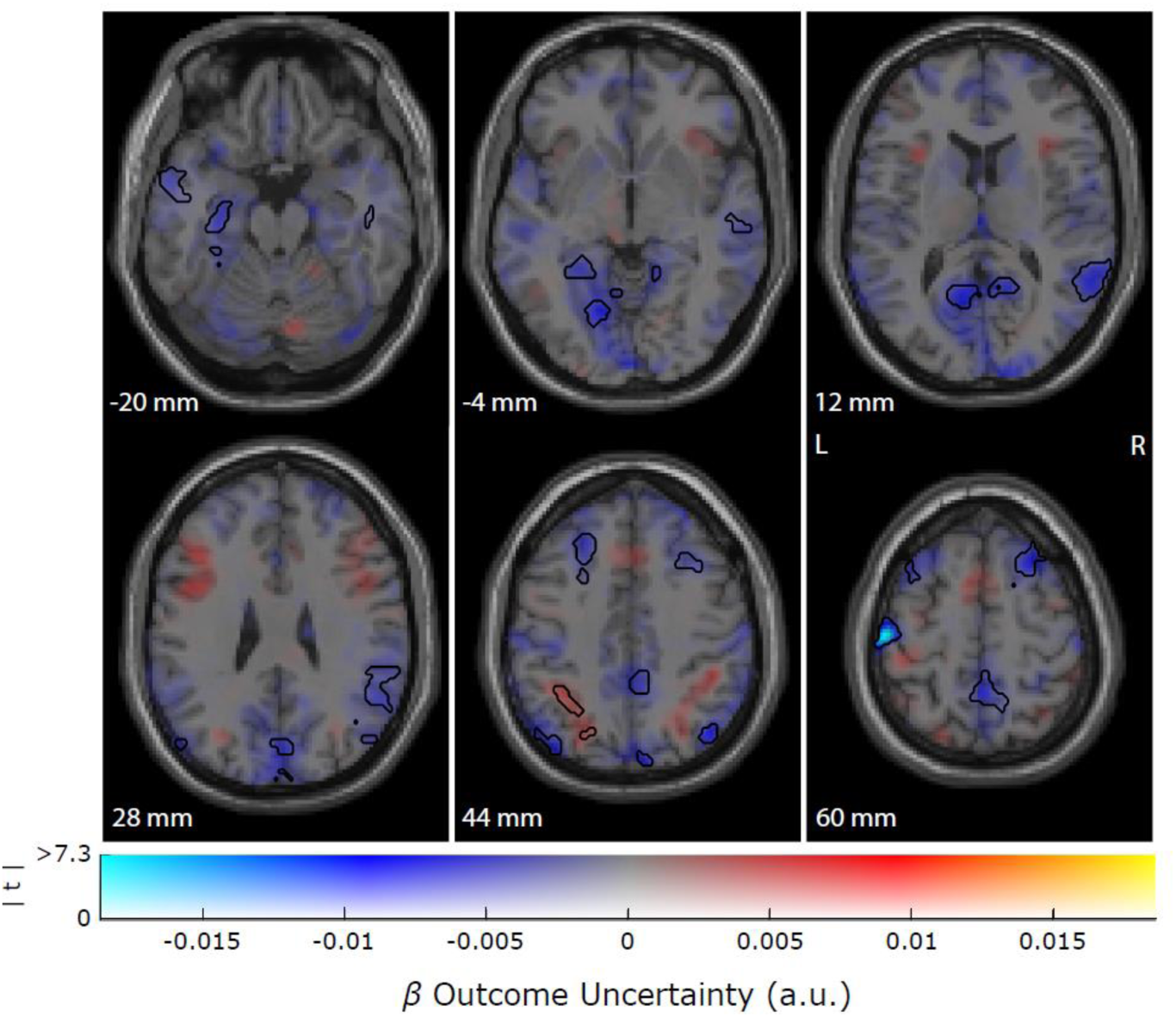
Clusters showing a positive (red) or negative (blue) activity as a function of OU. The maps show unthresholded values for OU and significant clusters (cluster-level corrected, FWE, *p* < .05) are encircled in black. The Z-coordinates correspond to the standard Montreal Neurological Institute (MNI) brain. Neuroimaging data are plotted using procedures introduced by Allen et al. (2012) and Zandbelt (2017).

**Table 1.**
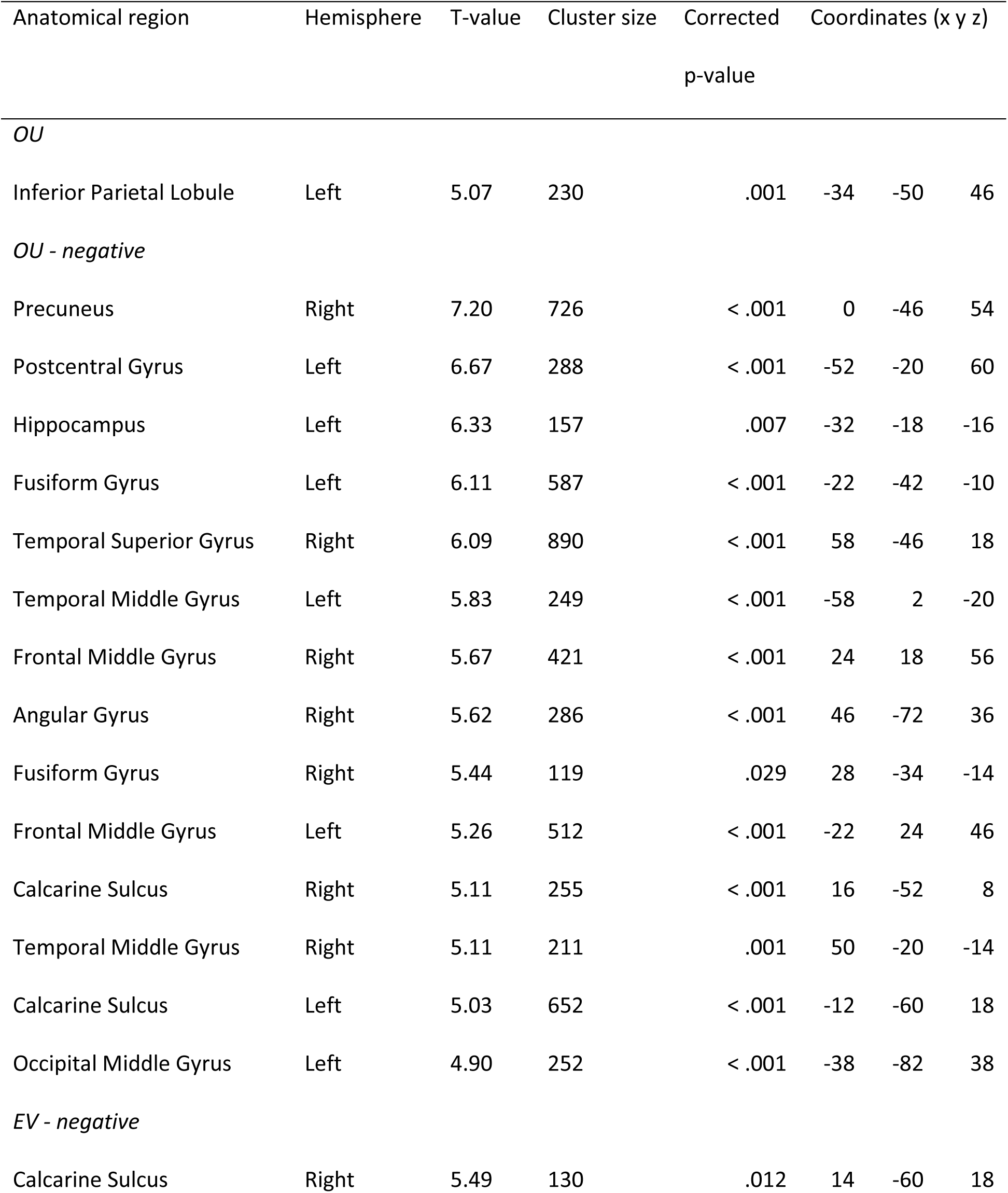
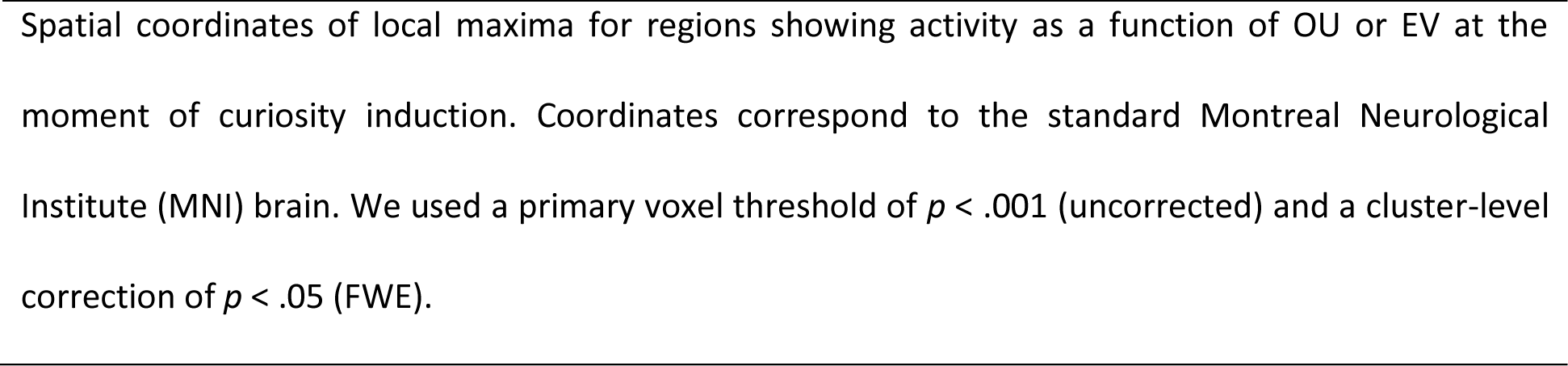
Brain regions associated with OU and EV during curiosity induction

| Anatomical region | Hemisphere | T-value | Cluster size | Corrected<br>p-value | Coordinates (x y z) |  |  |
| --- | --- | --- | --- | --- | --- | --- | --- |
| *OU* |  |  |  |  |  |  |  |
| Inferior Parietal Lobule | Left | 5.07 | 230 | .001 | -34 | -50 | 46 |
| *OU - negative* |  |  |  |  |  |  |  |
| Precuneus | Right | 7.20 | 726 | < .001 | 0 | -46 | 54 |
| Postcentral Gyrus | Left | 6.67 | 288 | < .001 | -52 | -20 | 60 |
| Hippocampus | Left | 6.33 | 157 | .007 | -32 | -18 | -16 |
| Fusiform Gyrus | Left | 6.11 | 587 | < .001 | -22 | -42 | -10 |
| Temporal Superior Gyrus | Right | 6.09 | 890 | < .001 | 58 | -46 | 18 |
| Temporal Middle Gyrus | Left | 5.83 | 249 | < .001 | -58 | 2 | -20 |
| Frontal Middle Gyrus | Right | 5.67 | 421 | < .001 | 24 | 18 | 56 |
| Angular Gyrus | Right | 5.62 | 286 | < .001 | 46 | -72 | 36 |
| Fusiform Gyrus | Right | 5.44 | 119 | .029 | 28 | -34 | -14 |
| Frontal Middle Gyrus | Left | 5.26 | 512 | < .001 | -22 | 24 | 46 |
| Calcarine Sulcus | Right | 5.11 | 255 | < .001 | 16 | -52 | 8 |
| Temporal Middle Gyrus | Right | 5.11 | 211 | .001 | 50 | -20 | -14 |
| Calcarine Sulcus | Left | 5.03 | 652 | < .001 | -12 | -60 | 18 |
| Occipital Middle Gyrus | Left | 4.90 | 252 | < .001 | -38 | -82 | 38 |
| *EV - negative* |  |  |  |  |  |  |  |
| Calcarine Sulcus | Right | 5.49 | 130 | .012 | 14 | -60 | 18 |

Spatial coordinates of local maxima for regions showing activity as a function of OU or EV at the moment of curiosity induction. Coordinates correspond to the standard Montreal Neurological Institute (MNI) brain. We used a primary voxel threshold of $p < .001$ (uncorrected) and a cluster-level correction of $p < .05$ (FWE).

ROI analyses in the ventral striatum showed no significant parametric modulation effects of OU (*t*(23) = -1.44, *p* =. 16) and EV (*t*(23) = -1.75, *p* = .093) at the time of curiosity induction.

*Curiosity Relief* - Next, we looked at the moment when participants’ curiosity was either relieved or not (i.e. when they were presented with the outcome or not). Three brain regions showed larger response when curiosity was relieved (Curiosity Relief – Yes > Curiosity Relief – No): the right insula, the right middle orbitofrontal cortex and the right inferior parietal lobule (Table 2, see Appendix; Figure 4). When the outcome was withheld (Curiosity Relief – No > Curiosity Relief – Yes), this led to larger response in the right middle occipital gyrus, the left frontal superior gyrus and the right frontal middle gyrus (Table 2, see appendix; Figure 4).

**Figure 4.**
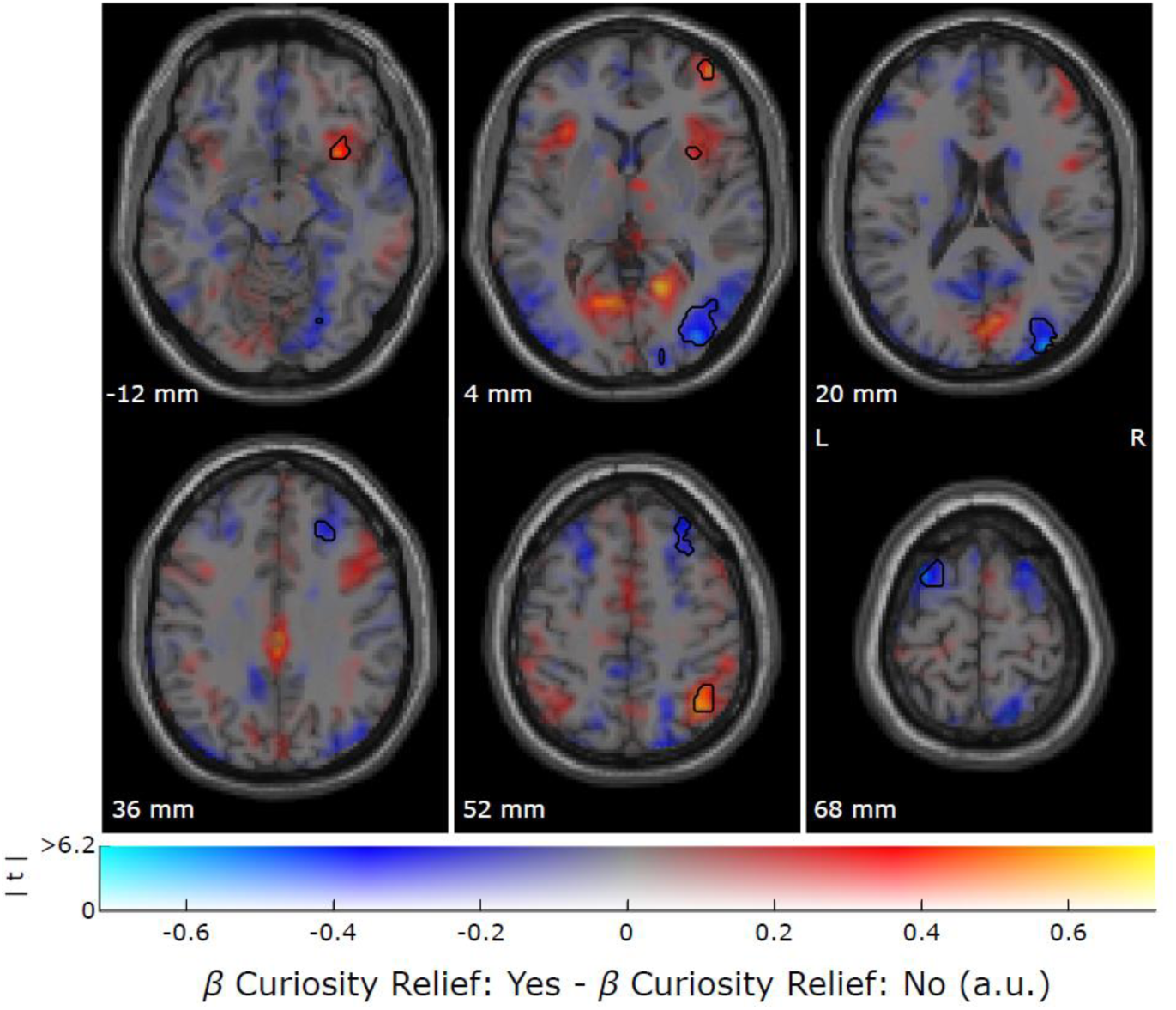
Clusters showing a positive (red) or negative (blue) activity when comparing relieved vs. non-relieved curiosity, at the time of the outcome presentation. The maps show unthresholded values for the curiosity relief contrast and significant clusters (cluster-level corrected, FWE, *p* < .05) are encircled in black. The Z-coordinates correspond to the standard Montreal Neurological Institute (MNI) brain. Neuroimaging data are plotted using procedures introduced by Allen et al. (2012) and Zandbelt (2017).

**Table 2.** Brain regions associated with curiosity relief

| Anatomical region | Hemisphere | T-value | Cluster size | Corrected<br>p-value | Coordinates (x y z) |  |  |
| --- | --- | --- | --- | --- | --- | --- | --- |
| *Curiosity Relief – Yes > Curiosity Relief – No* |  |  |  |  |  |  |  |
| Insula | Right | 6.14 | 124 | .018 | 36 | 12 | -12 |
| Middle orbitofrontal cortex | Right | 5.96 | 118 | .023 | 42 | 56 | 0 |
| Inferior parietal lobule | Right | 5.82 | 204 | .001 | 40 | -54 | 48 |
| *Curiosity Relief – No > Curiosity Relief – Yes* |  |  |  |  |  |  |  |
| Frontal superior gyrus | Left | 6.14 | 161 | .005 | -22 | 12 | 66 |
| Middle occipital gyrus | Right | 6.05 | 1076 | < .001 | 36 | -74 | 2 |
| Frontal middle gyrus | Right | 5.19 | 298 | < .001 | 30 | 34 | 48 |

Spatial coordinates of local maxima for regions showing activity as a function of outcome presentation (or not) at the moment of curiosity relief. Coordinates correspond to the standard Montreal Neurological Institute (MNI) brain. We used a primary voxel threshold of $p < .001$ (uncorrected) and a cluster-level correction of $p < .05$ (FWE).

We next investigated whether activity for nodes in these networks was linearly modulated by the size of the information update (i.e. IPE_relief_ and IPE_no relief_). Within the nodes that were activated when the outcome was presented and curiosity was relieved, the ROI analyses showed a linearly increasing response as a function of IPE_relief_ magnitude in the right insula (*t*(23) = 2.70, *p* = .013). This indicates that this region was involved in processing the prediction error of information when participants observed the outcome of a trial. No significant parametric modulation effects for IPE_relief_ were found in the right middle orbitofrontal cortex region (*t*(23) = -1.30, *p* = .21) or in the inferior parietal lobule region (*t*(23) = -0.59, *p*= .56). For the nodes that were active when the outcome was withheld, there was a significant parametric modulation effect of negative IPE (IPE_no relief_) in the right middle occipital gyrus (*t*(23) = -2.91, *p* = .0079), such that this area showed a higher activity on trials where participants were most uncertain about the outcome (highest OU, resulting in the largest negative IPE). The left frontal superior gyrus (*t*(23) = 0.96, *p* = .35), and the right frontal middle gyrus (*t*(23) = 0.91, *p* = .37), were not modulated by IPE_no relief_.

To summarize, the ROI analyses showed that the right insula was more active for higher positive information prediction errors (IPE_relief_), whereas the right middle occipital gyrus was more active for higher negative information prediction errors (IPE_no relief_). On the contrary, there was no significant parametric modulation effect of IPE_no relief_ in the right insula (*t*(23) = -1.30, *p* = .21) and for IPE_relief_ in the right middle occipital gyrus (*t*(23) = -1.66, *p* = .11). This indicates that the insula solely responded to prediction errors of information when the outcome was presented to the participants, such that this region was more active when they received a larger update of information. Likewise, the right middle occipital region solely responded to prediction errors of information when the outcome was not presented, such that this region was more active when uncertainty was highest, but curiosity was not relieved.

No significant parametric modulation of BOLD activity due to RPE (*t*(23) = 1.48, *p* = .15), IPE_relief_ (*t*(23) = -0.94, *p* = .36) or IPE_no relief_ (*t*(23) = -1.66, *p* = .11) were found in the ventral striatum at the time of curiosity relief.

## Discussion

In the current study, we examined the cognitive and neural mechanisms of curiosity. We found that curiosity and willingness to wait increased with increasing uncertainty about the outcome of a lottery. Furthermore, the curiosity induction through outcome uncertainty was related to increased BOLD activity in the inferior parietal lobule, whereas curiosity relief elicited increased BOLD activity in the right insula, orbitofrontal cortex and inferior parietal cortex. Importantly, outcome-related signal in the insula increased linearly with increasing information prediction error provided by the outcome (which was greatest when outcome uncertainty and thus curiosity was largest). Conversely, curiosity was not strongly related to the expected value of the lotteries.

## Curiosity is modulated by outcome uncertainty

Experiment 1 and 2 revealed that, even though information did not help participants to perform better or to obtain higher rewards, individuals were more curious about outcomes that provided them with a larger update of information (i.e. when outcome uncertainty was higher). Experiment 1 goes beyond previous studies on curiosity that have shown that humans and animals choose to seek information in order to reduce their uncertainty (i.e. Berlyne, 1962; Bromberg-Martin & Hikosaka, 2009, 2011; Lieberman, 1997), by showing that even in a passive observing task, humans show a drive towards gaining information and reducing uncertainty about what will happen.

One might argue that the findings of Experiment 1 might merely reflect participants’ compliance with task demands. Participants might have responded to be more curious about uncertain outcomes, simply because they felt that this was expected of them. To minimize such demand characteristics, we performed a second experiment (Experiment 2) in which curiosity was implicitly probed by means of participants’ willingness to wait for presentation of the outcome. We found that participants’ willingness to wait also increased with increasing outcome uncertainty. The finding that individuals are willing to “pay” with time for information to satisfy their curiosity supports the findings of Experiment 1 and makes is less likely that demand characteristics explain our results. Critically, this finding also suggests that a preference for information is present even when receiving advance information is costly (cf Roper et al., 1999) and when rewards have to be sacrificed (Blanchard et al., 2015; Stagner et al., 2010). This indicates that information gain may be intrinsically valuable and strengthens the hypothesis that it is evolutionarily adaptive to reduce our uncertainty about the state of the world (Gottlieb et al., 2013; Loewenstein, 1994).

Unlike the present study, prior work on human curiosity has suggested that individuals are mostly curious about information of intermediate uncertainty. For example, Kang et al (2009) have found that individuals are mostly curious about trivia questions about which they exhibit intermediate levels of confidence compared with questions about which they were very confident or not confident at all. This contrasts with our observation of a linear relationship between uncertainty and curiosity. However, this apparent discrepancy between prior work with trivia questions and our current work might reflect the possibility that participants in that prior work were not curious about questions with the highest uncertainty (i.e. the lowest confidence), because they were simply not interested in the topics of these questions. In other words, there may well be a correlation between one’s knowledge about a subject and one’s curiosity about the subject, which confounds the relationship between curiosity and uncertainty. In the present study, this confound was avoided by experimentally manipulating outcome uncertainty in a quantitative and controlled manner.

Somewhat surprisingly, we found no effects of expected value on curiosity. Perhaps our reward manipulation was not effective and rewards did not yield sufficient interest, because individuals had no way of maximizing their rewards. Alternatively, or additionally, the expected value computation might have been too demanding in the behavioral experiments. In the fMRI experiment we used only two types of vase configurations, which made it easier for the participant to encode both the outcome uncertainty and the expected value of the outcomes. Indeed, in this fMRI experiment we did find a significant increase in curiosity with increasing expected value. Nevertheless, this relationship was markedly smaller than the relationship between outcome uncertainty and curiosity, which remained robustly present after accounting for the link with expected value. As such, this finding strengthens the results from the behavioral studies, which demonstrate that uncertainty is a strong inducer of curiosity, over and above reward.

## Curiosity induction elicits parietal signal, whereas curiosity relief elicits frontal signal

Investigation of the neural BOLD signal during curiosity induction revealed a usual suspect as a function of outcome uncertainty, namely the parietal cortex (Huettel et al., 2005). Prior work has implicated this region in processing outcome uncertainty, and as such, our finding is unsurprising. The key novel finding is that BOLD signal in the insula covaried with the information prediction error at the moment of curiosity relief, such that activity in the insula linearly increased with the amount of information participants gained by seeing the outcome (information prediction error). Intriguingly, insular signals have previously been observed to vary with the risk prediction error, which represents the degree to which an outcome is surprising in the context of a gambling task (Preuschoff et al., 2008). In this study, risk prediction error signal was interpreted to reflect a role for the insula in risk prediction learning, relevant for learning when to avoid and when to approach risk. While risk- and information prediction errors are both measures of the degree to which outcomes are surprising in an uncertain environment, a key difference, however, is that the concept of risk is tightly linked to choice, whereas information prediction errors can be elicited in a task involving only the passive observation of events, as in the current study. As such, the current study suggest that the insula is involved not just in learning about risk in order to optimize choice, but may also have a role in knowledge acquisition by signaling information prediction error more generally.

In addition to these information prediction error signals in the right insula, we found overall increased BOLD signal the right orbitofrontal cortex and the right inferior parietal lobule at the time of curiosity relief. These findings generalize prior findings that the insula and orbitofrontal cortex respond to perceptual curiosity relief (Jepma et al., 2012) to non-perceptual curiosity relief, suggesting that these areas might be involved more generally in gaining information. As mentioned above, the insula might do so by computing specific information prediction error signals, which help us to optimize our current world model. The orbitofrontal cortex, however, has been argued to signal information about reward, reward-learning and regulation of reward-related cognition (Blanchard et al., 2015; Padoa-Schioppa, 2011; Rushworth et al., 2011; Wallis, 2007; Wilson et al., 2014). Given this established link between the orbitofrontal cortex and reward, it is perhaps unsurprising that this region was more active when participants saw the outcome than when they did not.

As mentioned above, we found that BOLD signal in the left inferior parietal lobule increased with increasing outcome uncertainty at the time of curiosity induction. This extends previous findings by Huettel et al. (2005), who found increased activity in parietal cortex with increasing outcome uncertainty in the context of a decision-making task. Given our finding that the inferior parietal lobule is similarly involved in processing outcome uncertainty in a passive observation task, it might be the case that this area is more generally involved in processing outcome uncertainty, which may in turn play a role in curiosity induction. This might be due to requirement of more attentive control in situations of high outcome uncertainty and, by consequence, high curiosity. Possibly, (curiosity-driven) actions associated with acquiring new information might require more attentive control, whereas actions with low uncertainty are more habitual and can be performed with little attention (Fan, 2014; Shenhav et al., 2013). Moreover, these curiosity-inducing signals in parietal cortex may operate independently from signals responsible for signaling expected rewards, as previous work in macaque monkeys (Foley et al., 2017) has indicated that parietal neurons may encode the expected reduction in uncertainty an action will bring, independent of expected reward and reward prediction errors.

## Conclusion

Our study sheds light on the cognitive and neural mechanisms of curiosity by demonstrating that outcome uncertainty is a strong driver of curiosity. This indicates that the more uncertain you are about the world around you, the more eager you will be to update your model of the world. This induction of curiosity is accompanied by BOLD signal in parietal cortex, whereas its release is accompanied by BOLD increases in the insula, orbitofrontal cortex and parietal cortex. Most strikingly, activity in the insula increased linearly with increasing information update provided by the outcome. Together, these findings point to a fundamentally adaptive role of curiosity as a drive to improve one’s current model of the world and implicate the parietal cortex and insula in this fundamental trait.

## Acknowledgements

This work was supported by The Netherlands Organisation for Scientific Research (NWO Vidi award 452-13-016 to FPdL and NWO Vici award 453-14-015 to RC), the James McDonell Foundation (JSMF scholar award 220020328 to RC) and the EC Horizon 2020 Program (ERC starting grant 678286 awarded to FPdL).

